# Cancer exosomes harbor diverse hypoxia-targeted mRNAs and contribute toward tumor angiogenesis

**DOI:** 10.1101/2020.06.12.147850

**Authors:** Pan Zhang, Su Bin Lim, Kuan Jiang, Ti Weng Chew, Boon Chuan Low, Chwee Teck Lim

## Abstract

Exosomes are extracellular vesicles of endosome origin secreted by various cells. The exosomal cargo, especially proteins and microRNAs, have been extensively investigated for their roles in intercellular communication and as biomarkers for clinical applications. However, the understanding of types and functions of exosomal mRNAs remains limited. Here, we evaluated the mRNAs of 61 hypoxia-targeted genes in exosomes by quantitative reverse transcription PCR (RT-qPCR). Among these 61 mRNAs, 14.8% of them were detected in the MCF10A-derived exosomes, 42.6% in the MCF7-derived exosomes, and 49.2% in the MDA-MB-231-derived exosomes, many of which are differentially regulated in response to hypoxic stress in a cell-line dependent manner. Consequently, 30 exosomal mRNAs are identified as cancer related biomarkers as they are present in cancer cell-derived exosomes and absent in MCF10A-derived exosomes. Co-culture of MDA-MB-231 cells with HUVECs shows uptake of MDA-MB-231 secreted exosomes by the Human umbilical vein endothelial cells (HUVECs). Subsequently, the cancer exosomal VEGFA mRNAs were translated within the HUVECs into proteins that promoted VEGFR-dependent angiogenesis. This finding provides novel insights into how cancer cells can directly contribute towards angiogenesis. RNA-seq also shows that cancer exosomes can upregulate epithelial-mesenchymal transition-related and metabolism-related genes. The transcripts of these genes are found present in the cancer exosomes, suggesting that the uptake of exosomal mRNAs at least partially contributed to the upregulation of the corresponding mRNAs. This study shows that cancer exosomes harbor diverse mRNAs, some of which can act as promising biomarkers as well as contribute towards reprogramming of recipient cells.

## Introduction

Exosomes are a type of extracellular vesicles with diameters between 30 nm and 150 nm^1^. These nano-sized, liquid bilayer-bound vesicles are secreted by cell upon fusion of multivesicular body with cell membrane^2^. Though the mechanism of packaging molecules into exosomes is not fully understood yet, the exosomes are known to deliver various functional proteins and nucleic acids, especially microRNAs (miRNAs), to recipient cells, which plays a critical role in intercellular communications^1,3^. Particularly, the tumor-derived exosomes are known to advance cancer progression by suppressing the immune response, educating the neighboring cells, and initiating the pre-metastatic niche formation^4^. Along with the growing understanding of exosome cargo, their clinical potential as cancer biomarkers has also triggered extensive investigations. Currently, certain exosomal proteins^5-8^, non-coding RNAs^9-13^ and mutated nucleic acids^14-16^ are proposed to be biomarkers for different types of cancer. However, prior studies have failed to evaluate the potential of the wild-type exosomal mRNAs as biomarkers, despite the presence of translatable wild-type mRNAs in the exosomes have been suggested.

Interestingly, a recent study demonstrated that some hypoxia-regulated mRNAs were present in the glioblastoma-derived exosomes and were modulated by hypoxic stimulus^17^. Hypoxia is a common signature of tumor microenvironment (TME) attributed to the insufficient supply of blood to the intensively proliferating tumor. In cancer cells, the hypoxic stress disturbs the oxygen-dependent degradation of hypoxia-inducible factors (HIFs), and thus, the preserved HIFs serve to transactivate various genes involved in angiogenesis, metastasis and resistance to therapy^18^. Consequently, hypoxia educates the TME, enhances tumor progression, and correlates with the poor prognosis of cancer patients^19^. As for hypoxia-induced exosomes, certain exosomal proteins^20-23^ and miRNAs^21,24-28^ have been identified in previous investigations to have contributed to enhanced angiogenesis, metastasis and immunosuppression, but the role of exosomal mRNAs has yet to be investigated.

Here, we investigated a panel of 61 hypoxia-targeted mRNAs in cancer cell-derived and normal cell-derived exosomes. We found that cancer cells shed 2.6-6.5 folds more exosomes per unit time than normal cells, and the size of MDA-MB-231-derived exosomes were larger than those from MCF10A and MCF7. Among these 61 hypoxia-targeted mRNAs, 14.8% of them were detected in the MCF10A-derived exosomes, 42.6% in the MCF7-derived exosomes, and 49.2% in the MDA-MB-231-derived exosomes. Consequently, 30 exosomal mRNAs were identified as cancer biomarkers as they were present in cancer cell-derived exosomes but absent in MCF10A-derived exosomes. Notably, these exosomal mRNAs were responsive to the hypoxic stress in a cell line-dependent manner. For MCF7, 92.3% of the altered mRNAs were downregulated, while for MDA-MB-231, 75% of the altered mRNAs were upregulated. Consistently, all of the 26 mRNAs that were present in exosomes derived from both cell lines showed remarkably higher levels in the hypoxia-induced MDA-MB-231-derived exosomes than hypoxia-induced MCF7-derived exosomes. Specifically, we showed that the VEGFA mRNAs were carried by exosomes derived from various types of cancer, transferred to HUVECs via exosomes, translated into proteins in HUVECs and eventually resulted in increase in VEGF-dependent angiogenesis. In addition, the hypoxia-induced MDA-MB-231-derived exosomes delivered the epithelial-mesenchymal transition (EMT)-related and the metabolism-related mRNAs to HUVECs, which possibly contributed to the elevation of these mRNAs in HUVECs.

## Results

### Quantitative analysis of the cell line-dependent exosome-generation rate

Exosomes were produced with incubation of three breast cell lines in serum-free medium - cancer cell MCF7 and MDA-MB-231 (MDA231), as well as non-tumorigenic cell MCF10A (Fig. 1A, left panel). With the lipophilic dye PKH26 that uniformly and thoroughly stained the cytoplasmic membranes of the cells, the exosomes isolated from the culture medium of the PKH26^+^ cells were also observed to have the PKH26 fluorescence (Fig. 1A, right panel). By western blotting, the enriched exosome-like vesicles from the culture medium were confirmed positive with exosome protein markers including Alix, CD63, CD9 and CD81 (Fig. 1B). With larger vesicles in the culture medium removed by 10,000 x g centrifugation for 30 min, the remaining vesicles were characterized by Nanoparticle Tracking Analysis (NTA) (Fig. 1C). The size of these vesicles ranged from 30 nm to 150 nm, which was consistent with that of exosomes^1^ (Fig. 1D, upper panel). These vesicles could be completely removed from the supernatant by 100,000 x g ultracentrifugation for 2 h, while exosomes were usually pelleted by the same condition^29^ (Fig. 1C). These results provided evidence that the vesicles obtained were exosomes.

**Fig. 1:**
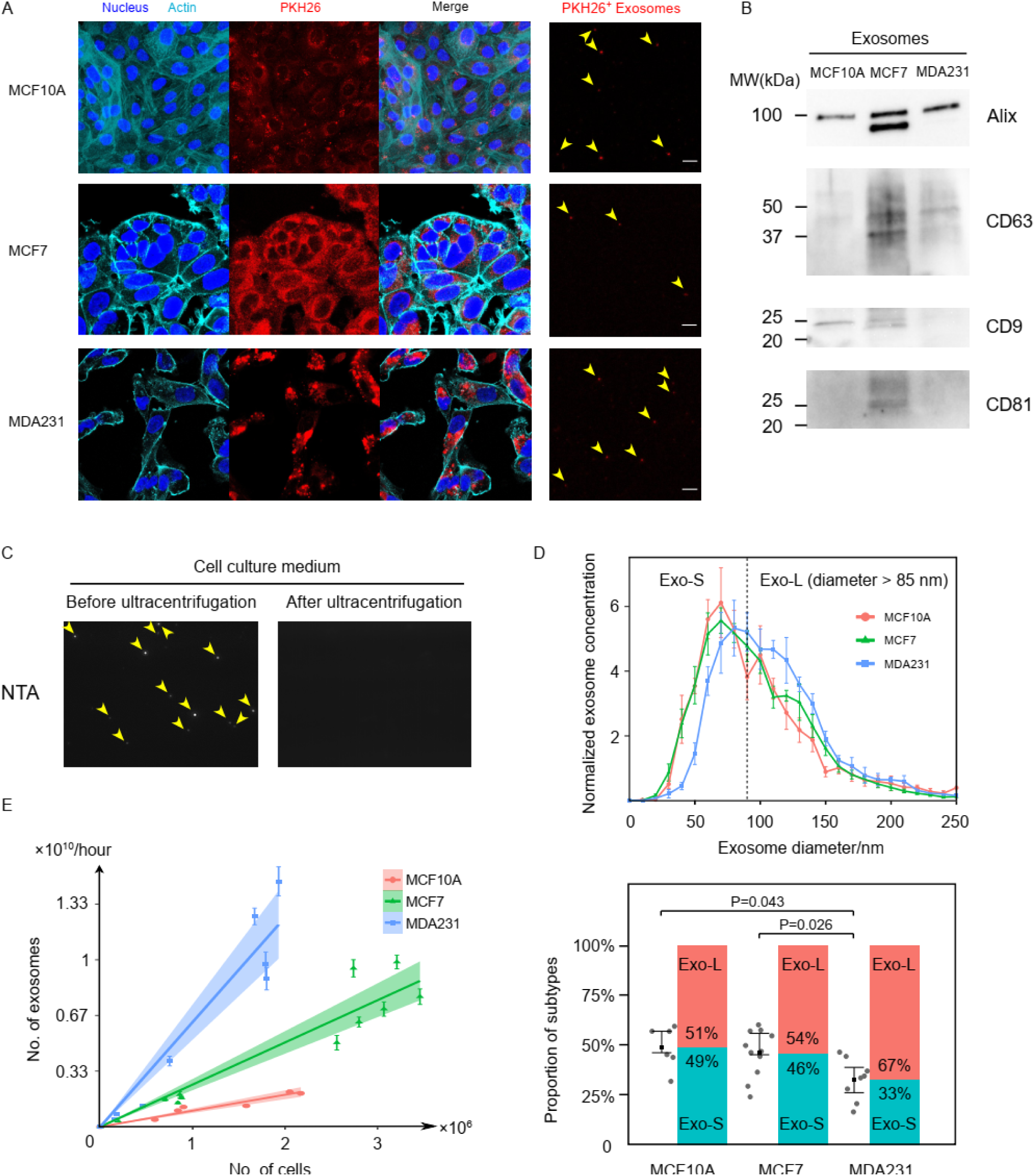
Characterization of cancer and normal cell derived exosomes. (A) Fluorescent imaging of MCF10A, MCF7 and MDA231 cells stained with PKH26. PKH26^+^ exosomes were identified in the culture medium of these PKH26^+^ cells. Scale bars = 5 µm. (B) Western blot result of Alix, CD63, CD9 and CD81 proteins in the isolated exosomes derived from MCF10A, MCF7 and MDA231. (C) Crude exosomes in the culture medium were assessed by the Nanoparticle Tracking Analysis. Ultracentrifugation was sufficient to remove all the exosomes in the cell culture medium. (D) Normalized size distribution of exosome for MCF10A (n=6), MCF7 (n=11) and MDA231 (n=8). Exo-S and Exo-L subtypes were separated by a diameter of 85 nm. Proportions of the two subtypes were calculated, with error bars indicating the 25th and 75th percentile. P values are calculated with Mann Whitney U test. (E) Scatter plot of exosomes number vs. cell number. Error bar of each point was generated by NTA. The data were fitted by linear regression with zero intercept. The highlighted area showed the 95% confidence interval of the coefficient.

The normalized size distribution of exosomes was averaged from six samples for MCF10A, 11 samples for MCF7 and eight samples for MDA231. Notably, the curve for MDA231 derived exosomes (MDA231-EXOs) showed a shift towards larger size compared to MCF10A derived exosomes (MCF10A-EXOs) and MCF7 derived exosomes (MCF7-EXOs) (Fig. 1D, upper panel). Dividing the exosome population into the small (EXO-S) and the large (EXO-L) subtypes by a 85 nm size cut-off according to a previous study^30^, the proportion of the EXO-L population for MDA231-EXOs (67.4% ± 3.7%) was substantially higher than MCF7 (53.9% ± 3.6%) and MCF10A (50.9% ± 4.4%) (Fig. 1D, lower panel). These results indicate that the metastatic cancer cell MDA231 tend to generate larger exosomes than MCF10A and MCF7 cells.

To quantitatively evaluate the exosome-generation rate of each cell line, we engaged coulter counter and NTA to count the cells and the secreted exosomes, respectively. Multiple measures were implemented to enhance the accuracy of the measurements (more details in Methods). Importantly, we noticed that the cell generated exosomes at a constant rate independent of the cell density since the exosome number linearly correlated with the cell number. By linear regression with zero intercept, the exosome-generation rates were calculated to be 0.96 × 10^9^ (95% confidence interval [CI] = 0.82 × 10^9^, 1.09 × 10^9^) per million cells per hour for MCF10A, 2.53 × 10^9^ (95% CI = 2.19 × 10^9^, 2.86 × 10^9^) per million cells per hour for MCF7, and 6.27 × 10^9^ (95% CI = 5.22 × 10^9^, 7.33 × 10^9^) per million cells per hour for MDA231 (Fig. 1E). Therefore, the exosome-generation rate is cell line-dependent, and cancer cells secrete exosomes at folds higher rate than healthy cells.

### Transcripts of hypoxia-targeted genes prevail in cancer exosomes

To assess the exosomal mRNAs, we established a panel of 61 genes which were reported to be transactivated by hypoxia-inducible factors (HIFs) and contributed to tumor progression^31,32^. These genes are mainly involved in seven types of biological processes: angiogenesis (VEGFA, CXCL12, KITLG, PDGFB, PGF), extracellular matrix (ECM) stiffening (LOX, LOXL2, P4HA1, P4HA2, PLOD2), cell motility (MET, SNAIL1, SNAIL2, ROCK1, RHOA, AXL, AMF, TWIST, ZEB1, ZEB2, TCF3), metastasis (PLAUR, ANGPTL4, ANGPT2, MMP14, MMP2, MMP9, CXCR4, L1CAM), cancer stem cell maintenance (IL6, IL8, IL19, CD47, SOX2, TAZ, POU5F1, MYC, KLF4, NANOG, ALKBH5), metabolism (GLUT1, PLIN2, BNIP3, BNIP3L, SHMT2, LONP1, SLC16A4, PHGDH, PKD1, MXI1, SIAH2, LDHA, CA9, FABP3, FABP7, COX4I2, NDUFA4L2, NHE1), and immune evasion (NT5E, CD39, PDL1) (Supplementary Table 1; Fig. 2A).

**Fig. 2:**
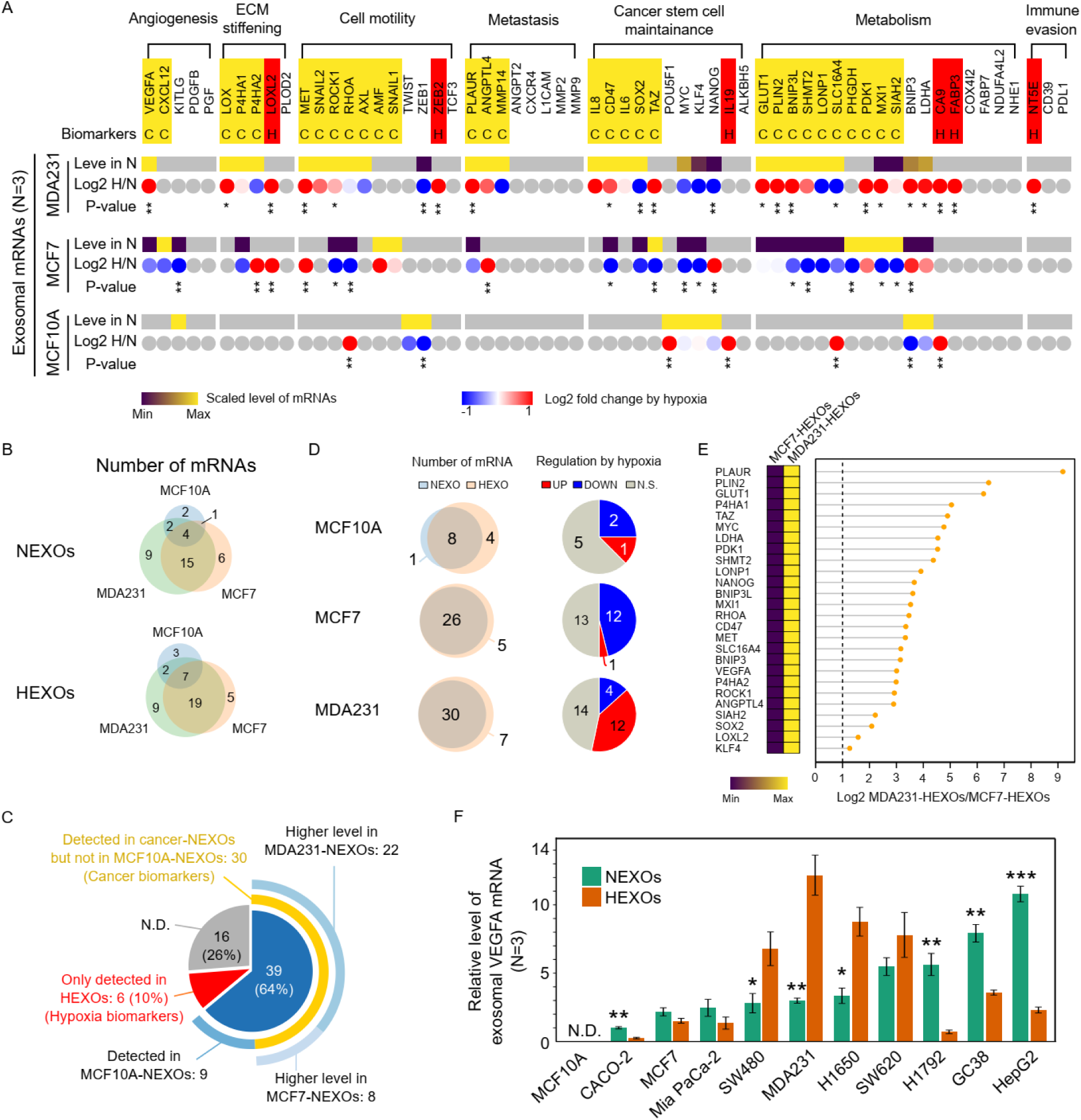
Quantification of hypoxia-targeted mRNAs in exosomes. (A) The heatmap with square elements presents exosomal mRNA levels relative to exosomal GAPDH and scaled by column. The heatmap with round elements shows log2 fold change of mRNA level by the hypoxic stimulus. Each expression level is the average of three independent experiments. (B) Venn diagram shows the number of mRNAs present in exosomes derived from each cell lines. (C) Pie chart of the 61 hypoxia-targeted mRNAs classified by their presence in different exosomes. (D) Venn diagram shows the number of mRNAs in normoxic and hypoxic exosomes. Pie chart presents the number of exosomal mRNAs significantly regulated by hypoxia. (E) All of the 26 mRNAs shared by MCF7-HEXOs and MDA231-HEXOs exhibited higher levels in MDA231-HEXOs. (F) The levels of VEGFA mRNAs in the exosomes derived from multiple cancer cell lines, with or without the hypoxic stress. *P < 0.05, **P < 0.01. ***P < 0.001 based on student’s t test for (A) and (F).

To enable the quantification of exosomal mRNA by RT-qPCR, the housekeeping gene GAPDH was verified to be ubiquitously present in exosomes, and thus used for normalization of expression levels (Fig. S2A). By RT-qPCR, the expressions of a small list of ten mRNA expressions were first quantified and compared between cell and exosome (Fig. S2B). Notably, the genes MET, P4HA2, ANGPTL4 and VEGFA showed higher expression level in MCF10A cells than MDA231 cells. However, the MDA231 cells loaded these mRNAs into exosomes, while the MCF10A cells did not. Furthermore, the hypoxia-induced change of MET mRNA level in exosomes was in contrary to that in the cells (Fig. S2C). These inconsistencies of mRNA level between the exosomes and cells demonstrated that the two were not necessarily correlated.

Next, we quantified the exosomal mRNAs of the 61 hypoxia-targeted genes for MCF10A, MCF7 and MDA231 with or without the hypoxic stimulus (Supplementary Data 3). The coefficient of variance of the quantifications were all less than 0.3, suggesting that exosomal mRNA levels were homeostatic and the quantification of exosomal mRNA by RT-qPCR was reproducible (Fig. S2D). As a result, 73.8% (45/61) of the transcripts were detected in at least one group of the exosomes (Fig. 2A, C). Remarkably, the cancer exosomes carried substantially more types of mRNAs than normal exosomes under either normoxia or hypoxia, with 14.8% (9/61) mRNAs in normoxic MCF10A-EXOs (MCF10A-NEXOs), 42.6% (26/61) mRNAs in normoxic MCF7-EXOs (MCF7-NEXOs), 49.2% (30/61) mRNAs in normoxic MDA231-EXOs (MDA231-NEXOs), 19.7% (12/61) mRNAs in hypoxic MCF10A-EXOs (MCF10A-HEXOs), 50.8% (31/61) mRNAs in hypoxic MCF7-EXOs (MCF7-HEXOs), and 60.7% (37/61) mRNAs in hypoxic MDA231-EXOs (MDA231-HEXOs) (Fig. 2A, B).

Altogether, a total of 49.2% (30/61) mRNAs were found present in normoxic cancer exosomes but absent in MCF10A-NEXOs, highlighting their potential as cancer biomarkers. Among the 30 genes, 26.7% (8/30; CXCL12, AMF, SNAIL1, TAZ, PHGDH, PDK1, MXI1,SIAH2) showed higher level in MCF7-NEXOs and the other 73.3% (22/30; VEGFA, LOX,P4HA1, P4HA2, MET, SNAIL2, ROCK1, RHOA, AXL, PLAUR, ANGPTL4, MMP14, IL8, CD47, IL6, SOX2, GLUT1, PLIN2, BNIP3L, SHMT2, LONP1, SLC16A4) showed higher level in MDA231-NEXOs (Fig. 2A, C, S2E). The hypoxia-induced alteration in the exosomal mRNAs was assessed comparatively in the normoxic and hypoxic exosomes. We observed that more mRNAs were packed into exosomes under the hypoxic stress, with particularly four more mRNAs (RHOA, IL19, SLC16A4, CA9) for MCF10A-EXOs, five more mRNAs (P4HA2, LOXL2, MET, ANGTPL4, NANOG) for MCF7-EXOs, and seven more mRNAs (LOXL2, ZEB2, TAZ, PDK1, CA9, FABP3, NT5E) for MDA231-EXOs (Fig. 2A, D). Six of these hypoxia-induced exosomal mRNAs (LOXL2, ZEB2, IL19, CA9, FABP3, NT5E) were absent in any normoxic exosomes, which can possibly be applied to monitor the extent of hypoxic stress in the TME (Fig. 2A, S2F).

Notably, the mRNA levels in the MCF7-HEXOs were drastically distinguishable from that in the MDA231-HEXOs. There were 26 mRNAs present in both MCF7-HEXOs and MDA231-HEXOs, and these mRNAs all exhibited more than 2 folds higher levels in the MDA231-HEXOs than the MCF7-HEXOs (Fig. 2E). This result was consistent with the observation that MCF7 and MDA231 responded to hypoxia in an opposite manner in terms of the exosomal mRNA level. For MCF7-EXOs, the hypoxia induced significant changes of 13 mRNAs, with 92.3% (12/13) of them downregulated. Meanwhile for MDA231-EXOs, 16 mRNAs were prominently altered by hypoxia, with 75% (12/16) of them upregulated (Fig. 2D, right). These findings suggested that in the loading of mRNAs into exosomes, the hypoxic stress may be suppressive for benign cancer cells but favorable for aggressive cancer cells.

The ubiquity of the exosomal hypoxia-targeted mRNAs were verified for multiple cancer cell lines with the VEGFA selected as a candidate for proof-of-principle. Importantly, VEGFA mRNA was detected in exosomes from all eight cancer cell lines we tested, including colorectal cancer cells (CACO-2, SW480, SW620), pancreatic cancer cells (MiaPaCa-2), lung cancer cells (H1650, H1792), gastric cancer cells (GC38), and liver cancer cells (HepG2). Together with the breast cancer, 70% (7/10) of the tested cancer cells showed significant changes in the exosomal VEGFA mRNA level induced by hypoxia (Fig. 2F). Interestingly, the VEGFA protein levels in exosomes and cell lysates were not modulated by hypoxia, indicating that the impacts of hypoxia on the loading of mRNAs into exosomes can be different from that of proteins. (Fig. S2G).

### Uptake of exosomal RNAs by HUVECs

To test the hypothesis that cancer exosomes deliver mRNAs to recipient cells, we co-cultured MDA231 cells and HUVECs (Fig. 3A). To first observe the uptake of cancer exosomes by HUVECs, the CD63-GFP^+^ MDA231 cell line was established and stained with PKH26 before coculture with HUVECs, so that the MDA231-EXOs were likely to be labeled either by CD63 or PKH26. (Fig. 3B). After 24 h co-culture of CD63-GFP^+^ PKH26^+^ MDA231 and HUVECs, both PKH26 and CD63 were spotted inside some of the HUVECs with some of their spots overlapped, demonstrating that the CD63^+^ exosomes or the PKH26^+^ exosomes derived from MDA231 were internalized by HUVECs within the period of 24 hours (Fig. 3C). To further demonstrate the transfer of RNA from MDA231 to HUVECs, the two cell lines were cocultured, with the RNAs of the PKH26^+^ MDA231 stained by the SYTO RNAselect green fluorescent dye which was stable for more than 24 hours (Fig. 3D, E). Staining the PKH26^+^ MDA231-EXOs with the SYTO RNAselect dye showed the presence of RNAs in some of the exosomes (Fig. 3F). Live cell imaging of HUVECs co-cultured with the SYTO RNAselect^+^ PKH26^+^ MDA231 cells showed a gradual accumulation of fluorescent RNAs in HUVECs within 15 hours (Fig. 3G, H). These results demonstrated that MDA231 cells transferred RNAs to HUVECs via exosomes.

**Fig. 3:**
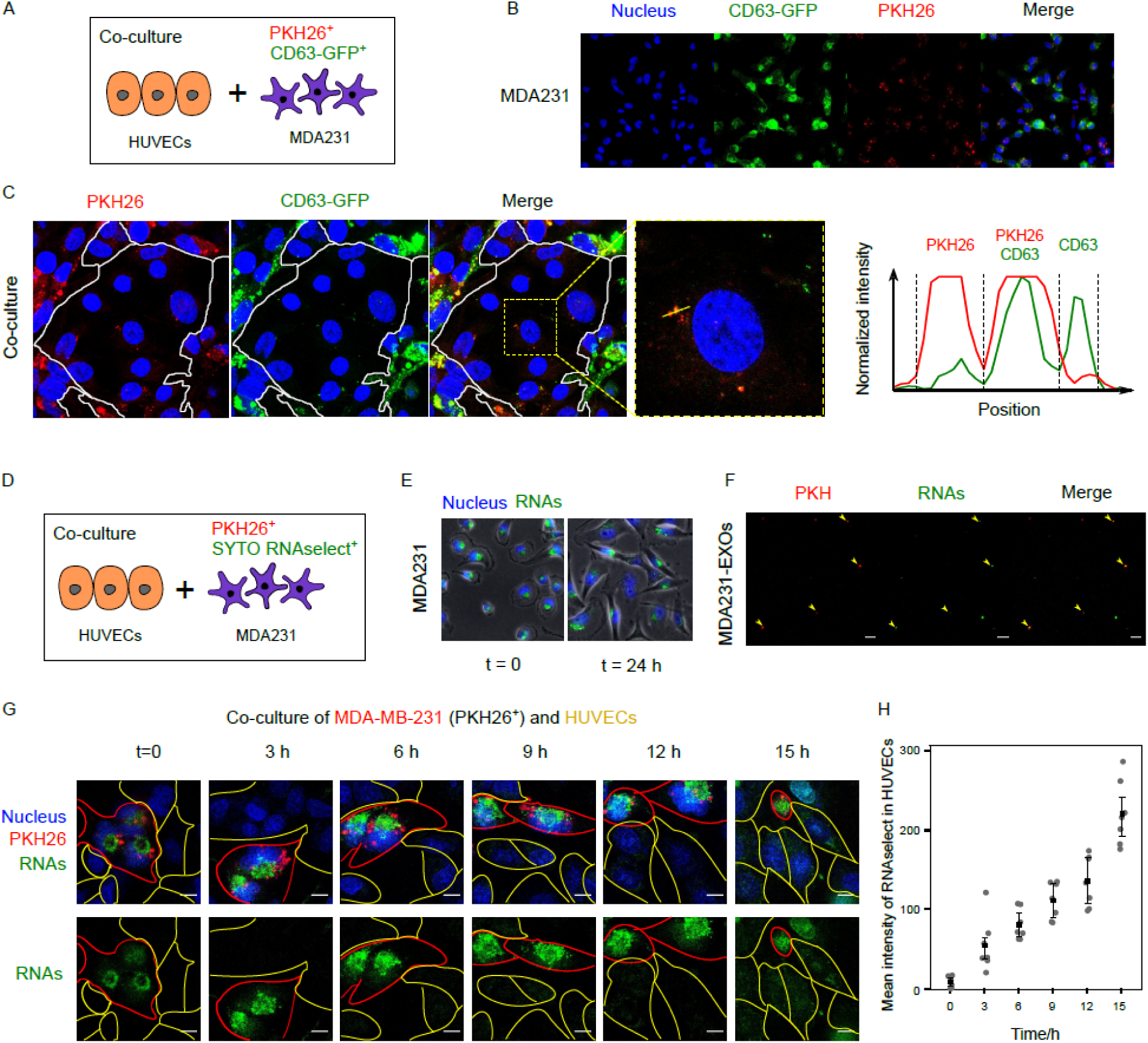
MDA231 transfers exosomes and exosomal RNAs to HUVECs. (A) HUVECs were co-cultured with CD63-GFP^+^ and PKH26^+^ MDA231. (B) MDA231 cells were transfected with CD63-GFP and stained by PKH26. (C) After co-culture for 24 hours, CD63-GFP^+^ and PKH26^+^ exosomes were internalized by HUVECs. Intensity plot shows the partial overlapping of CD63-GFP and PKH26. (D) HUVECs were co-cultured with SYTO RNASelect^+^ and PKH26^+^ MDA231. (E) The SYTO RNAselect specifically stained RNAs and lasted for more than 24 hours. (F) The SYTO RNAselect stained some of the PKH26^+^ exosomes. (G) Live cell imaging of co-culture. The stained RNAs from MDA231 gradually accumulated in HUVECs in 15 hours. Scale bars = 10 µm. (H) The average intensity of RNAs in multiple HUVECs was measured by ImageJ (n=7).

### Translation of exosomal VEGF mRNAs by HUVECs

To show that translatable mRNAs can be delivered by exosomes, we tagged the VEGF with GFP and tracked the transfer of VEGFA-GFP from MDA231 to HUVECs through exosomes (Fig. 4A). The VEGFA-GFP plasmid was constructed with the fusion of GFP at the c-terminus of VEGF165 so that the biological activity of VEGFA remained undisturbed^33^. With transient transfection, MDA231 cells expressed VEGFA-GFP and loaded an appreciable amount of VEGF-GFP mRNA but not VEGF-GFP protein into exosomes (Fig. 4B, D, E). Treated with the exosomes secreted by the VEGFA-GFP^+^ PKH26^+^ MDA231 for 24 hours, HUVECs were found to be positive with the GFP mRNA, VEGF-GFP protein and the PKH26 fluorescence (Fig. 4C, E). Therefore, translatable mRNAs can be delivered to recipient cells through exosomes.

**Fig. 4:**
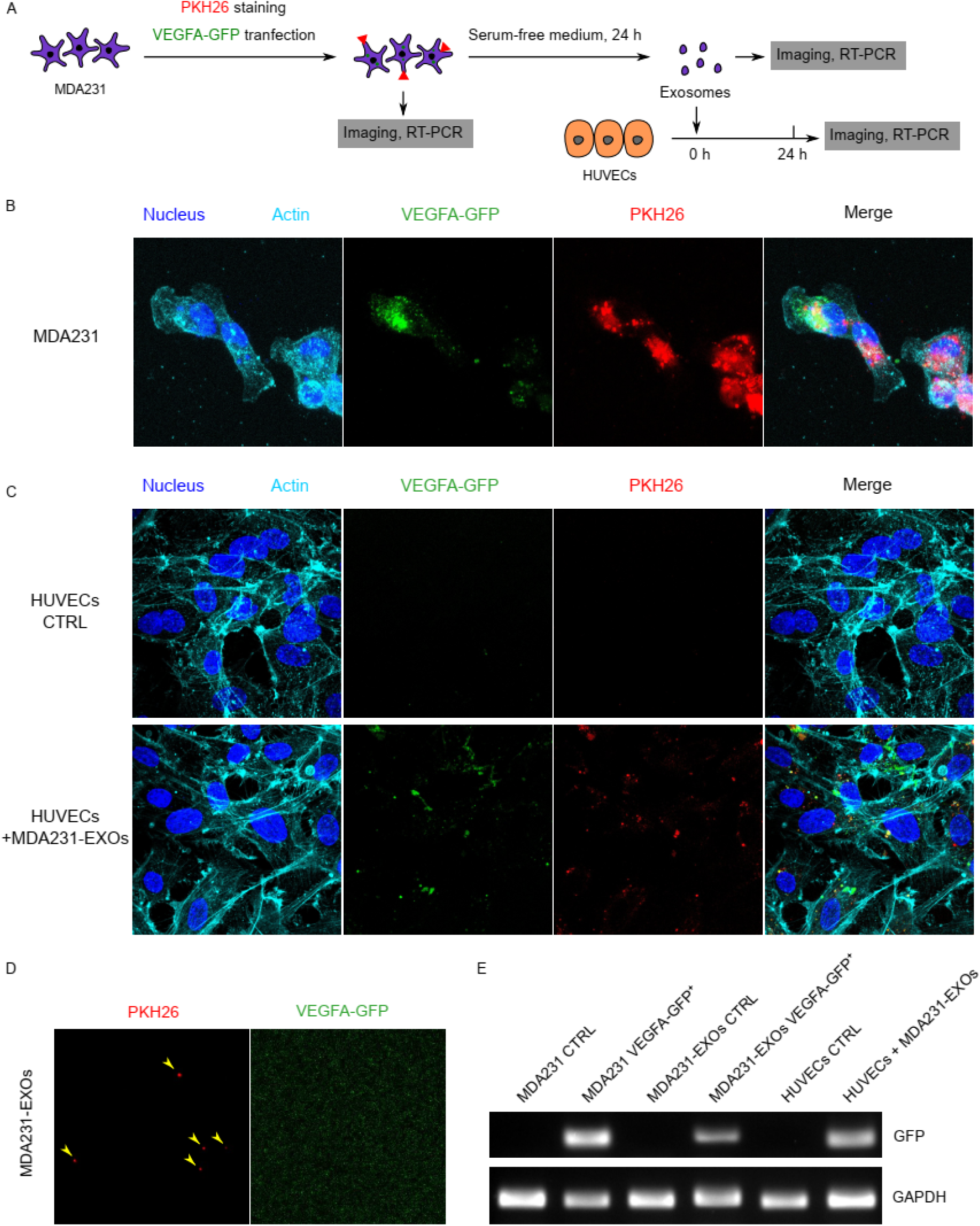
Exosomal VEGFA-GFP mRNA was internalized and translated by HUVECs. (A) The workflow of co-culture of HUVECs with exosomes derived from MDA231 transfected with VEGFA-GFP. (B) MDA231 were transfected with VEGFA-GFP and stained with PKH26. (C) With incubation of exosomes for 24 h, HUVECs internalized the PKH26^+^ exosomes that carried VEGFA-GFP mRNA and synthesized VEGFA-GFP protein. (D) The VEGFA-GFP protein was absent in PKH26^+^ exosomes derived from MDA231 transfected with VEGFA-GFP. (E) DNA gel electrophoresis proved that GFP mRNA was absent in controls and present in the MDA231 transfected with VEGFA-GFP, the exosomes derived from the transfected MDA231, and the HUVECs treated with the VEGFA-GFP mRNA^+^ exosomes.

### MDA231-HEXOs promoted VEGFR-dependent angiogenesis

The hypoxia-induced VEGF is recognized as a major driving force for angiogenesis^34^. We asked if the hypoxia-induced cancer exosomes that carried abundant VEGF mRNA contributed to angiogenesis. The tube formation assay of the HUVECs revealed that MDA231-HEXOs substantially increased the total length of tubes after 24 h. This increase was completely halted by the VEGF receptor inhibitor Axitinib (Fig. 5A, B). Using RT-qPCR, expressions of VEGFA, VEGFR1 and VEGFR2 which are the key genes for VEGF-dependent angiogenesis pathway, were found to be significantly upregulated in the HUVECs treated with the MDA231-HEXOs. On the other hand, the TIE-2 dependent angiogenesis pathway related key genes including ANGPT1, ANGTP2, TIE1 and TIE2, remained unaffected (Fig. 5C). Moreover, western blotting confirmed the enhanced phosphorylation of Erk1/2, which is a downstream of the angiogenic signaling transduction, and the elevation of the VEGFR1 protein (Fig. 5D).

**Fig. 5:**
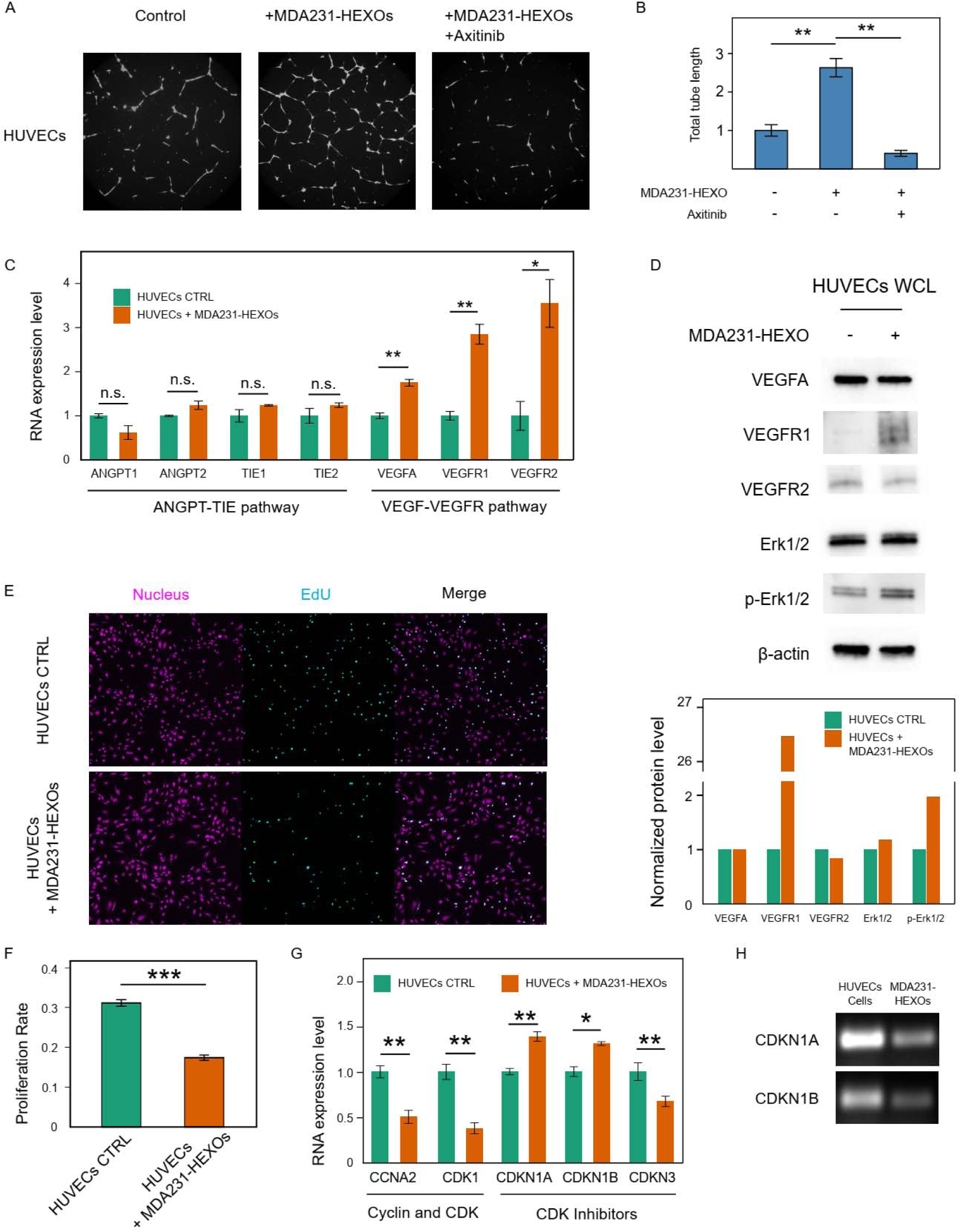
Treating cancer exosomes promotes VEGFR-dependent angiogenesis. (A) Tube formation assay of HUVECs in basal medium, treated with MDA231-HEXOs, or treated with both MDA231-HEXOs and Axitinib for 24 h. (B) Normalized total tube length analyzed by ImageJ (n=6). (C) Treatment with MDA231-HEXOs induced changes of gene expressions in HUVECs (n=3). (D) Western blot of proteins related to VEGF-dependent angiogenesis pathway. Average intensities of the bands were first normalized to β-actin, and then normalized to the control group for each protein. (E) EdU proliferation assay of HUVECs in basal medium or treated with MDA231-HEXOs for 24 h. (F) Proliferation rate of HUVECs were represented by the ratio of EdU^+^ cells to the Hoechst^+^ cells (n=25). (G) The expressions of proliferation-related genes that were significantly altered by the treatment of MDA231-HEXOs (n=3). (H) The two upregulated CDK inhibitors, CDKN1A and CDKN1B, were carried by MDA231-HEXOs. *P < 0.05, **P < 0.01. ***P < 0.001 based on Mann-Whitney U test for (B) and (F), or Student’s t test for (C) and (G). Error bars indicate the standard error of the mean (SEM) for (B)(C)(F) and (G).

Activation of angiogenesis pathway is known to boost cell proliferation. However, an EdU assay showed that the proliferation rate of HUVECs was reduced by the MDA231-HEXOs from 31.1% ± 0.83% to 17.4% ± 0.68% (Fig. 5E, F). This suppression of cell proliferation was contradictory to the prevalently reported pro-proliferative effect of cancer exosomes^15,35-39^. We assessed the expressions of six cyclins (CCNA1, CCNA2, CCND1, CCND2, CCND3, CCNE2), five cyclin-dependent kinase (CDK1, CDK2, CDK4, CDK6, CDK7) and eight cyclin-dependent kinase inhibitors (CDKN1A, CDKN1B, CDKN1C, CDKN2A, CDKN2B, CDKN2C, CDKN2D, CDKN3) in HUVECs. In line with the suppression of proliferation, the pro-proliferative CCNA2 and CDK1 were downregulated and the anti-proliferative CDKN1A and CDKN1B were upregulated with the treatment of MDA231-HEXOs (Fig. 5G). Notably, the mRNAs of the two upregulated genes CDKN1A and CDKN1B were carried by the MDA231-HEXOs (Fig. 5H).

### MDA231-HEXOs elevated the metabolism-related and motility-related mRNAs in HUVECs

The MDA231-HEXOs-mediated changes of HUVECs transcriptome were investigated using RNA sequencing. Two control samples and two exosome-treated samples were analyzed. The correlation assay demonstrated good reproducibility within the two groups (Fig. S3B). The sequencing reads were processed by multiple packages to minimize the algorithm-dependent bias (more details in Methods). Eventually, we acquired a list of differentially expressed genes (DEGs) comprising 566 genes, with 42.9% (243/566) DEGs upregulated and the rest 57.1% (323/566) DEGs downregulated by the treatment of MDA231-HEXOs (Fig. 6A, S3A, C; Supplementary Data 4). Prominently, the cell component enrichment analysis showed that 24.0% (136/566; 53 upregulated, 83 downregulated) of the DEGs were related to the extracellular exosomes which was the cause of these DEGs (Fig. 6B).

**Fig. 6:**
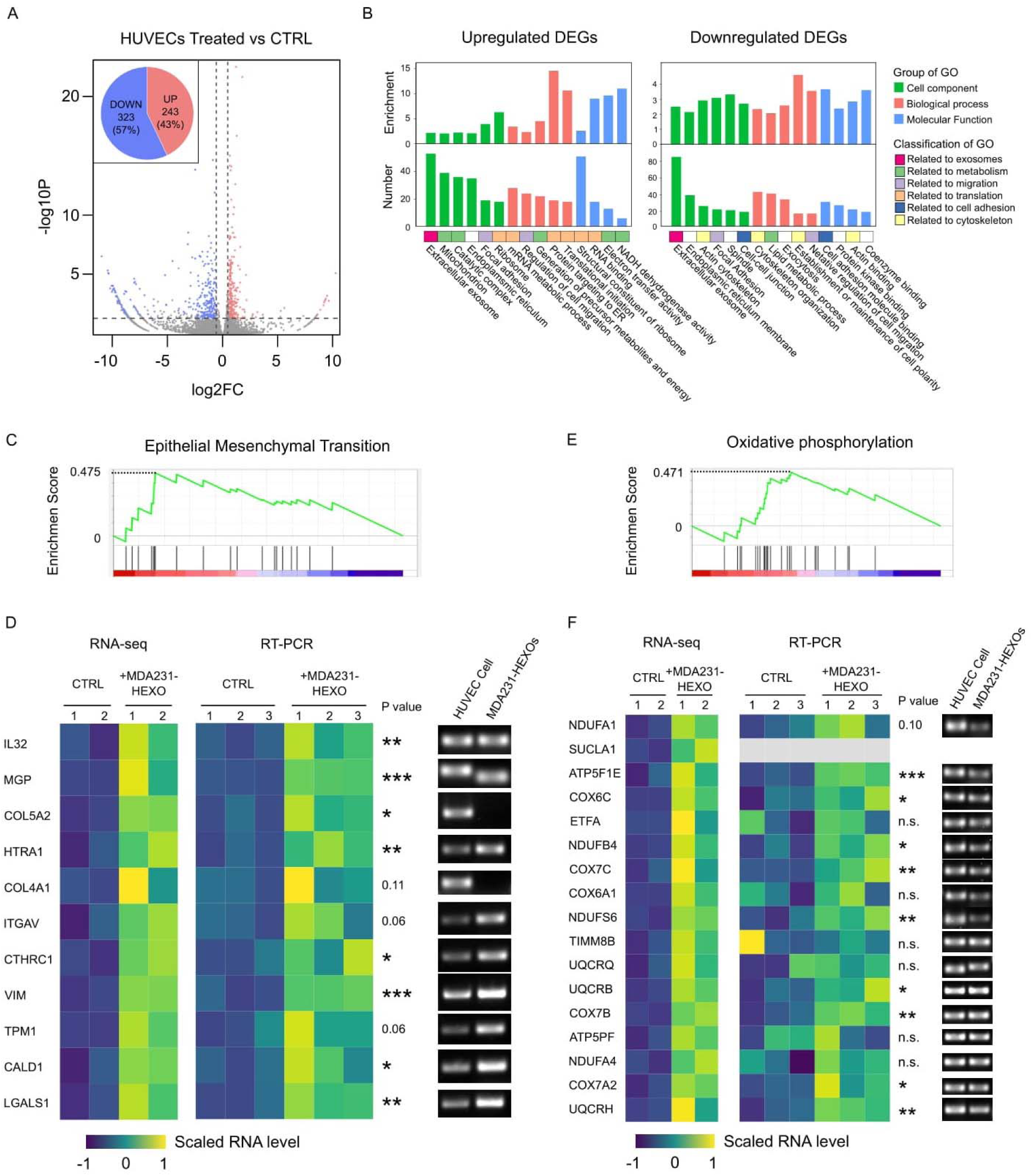
RNA-seq of HUVECs treated with cancer exosomes. (A) Volcano plot shows RNA-seq data of HUVECs with or without the treatment of MDA231-HEXOs that were analyzed with HTSeq and DESeq2 packages. Differentially expressed genes (DEGs) were highlighted with red for 242 upregulated genes and blue for 323 downregulated genes. (B) Gene ontology analysis of DEGs. Significantly enriched GO terms with large number of related genes were plotted. (C)(E) The EMT gene set and OP gene set were significantly enriched in the DEGs with 0.475 (P=0.017) and 0.471 (P=0.004) enrichment score, respectively. (D)(F) Expression levels of the upregulated DEGs that belong to the EMT gene set or OP gene set as quantified by RNA-seq or RT-qPCR. DNA gel electrophoresis demonstrated the presence of these DEGs in MDA231-HEXOs except COL5A2, COL4A1 and SUCLA1. *P < 0.05, **P < 0.01. ***P < 0.001 based on Student’s t test.

**Fig. 7:**
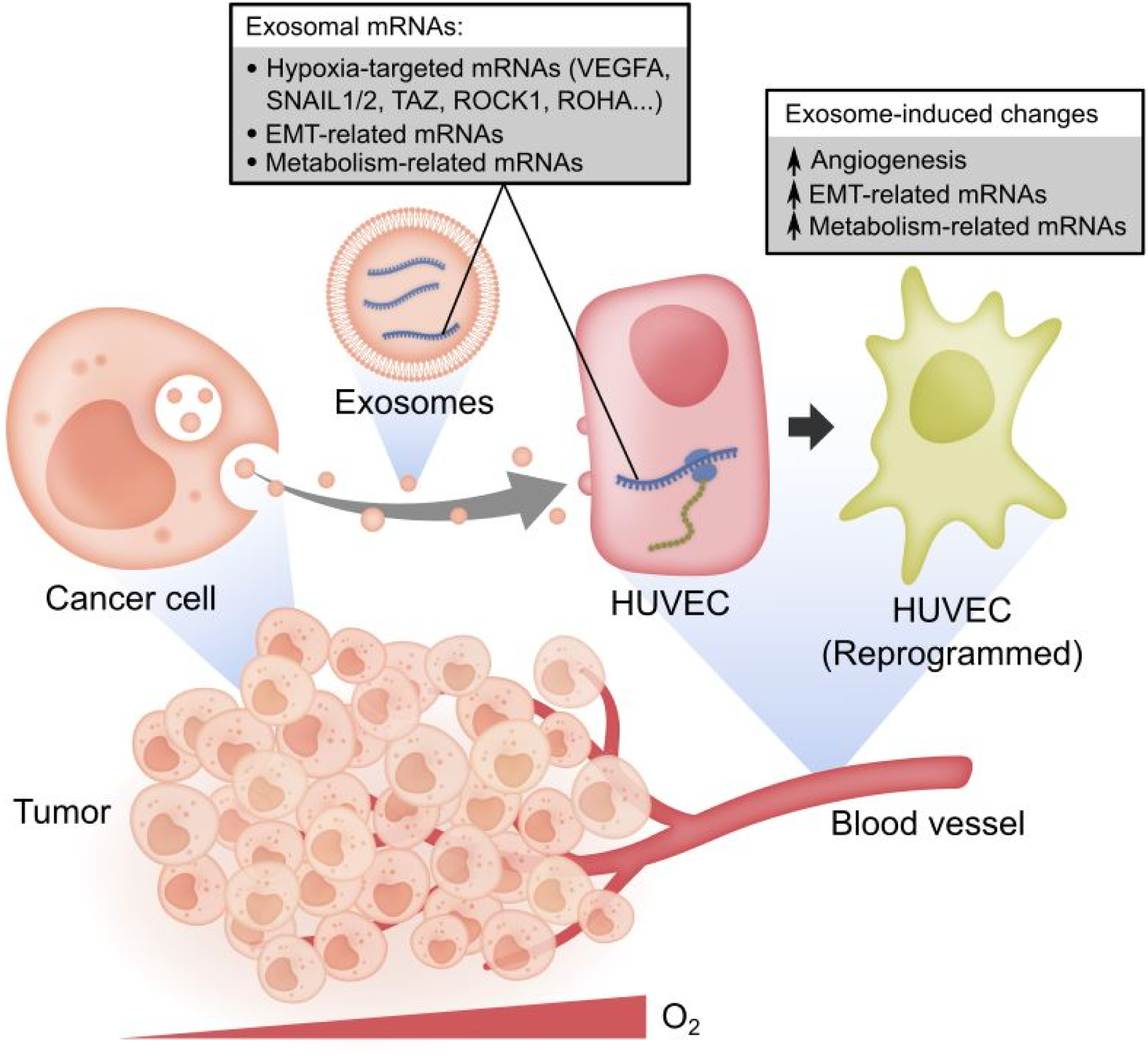
Summary schematic. Exosomes derived from tumor cells carried diverse mRNAs including hypoxia-targeted mRNAs, EMT-related mRNAs and metabolism-related mRNAs. Some of these exosomal mRNAs were responsive to the hypoxic stimulus in the tumor microenvironment. The tumor-derived exosomes delivered translatable mRNAs to HUVECs and reprogramed the HUVECs on promoting the angiogenesis and upregulating the EMT-related and metabolism-related gene expressions.

The gene ontology (GO) enrichment analysis of the DEGs revealed the pro-motility and pro-metabolism signatures of the DEGs. For the motility, the cell migration related genes were enriched in both upregulated and downregulated DEGs, and the cell adhesion related genes were enriched in the downregulated DEGs (Fig. 6B). For the metabolism, several related GO terms were enriched in the upregulated DEGs (Fig. 6B). Consistently, the gene set enrichment analysis (GSEA) identified two enriched hallmark gene sets: the EMT gene set (enrichment score [ES] = 0.475, P = 0.017) and the oxidative phosphorylation (OP) gene set (ES=0.471, P=0.004) (Fig. 6C, E).

Particularly, the EMT-related gene panel included 11 upregulated genes (IL32, MGP, COL5A2, HTRA1, COL4A1, ITGAV, CTHRC1, VIM, TPM1, CALD1, and LGALS1), and the OP-related gene panel involved 17 upregulated genes (NDUFA1, SUCLA1, ATP5F1E, COX6C, ETFA, NDUFB4, COX7C, COX6A1, NDUFS6, TIMM8B, UQCRQ, UQCRB, COX7B, ATP5PF, NDUFA4, COX7A2, UQCRH) (Fig. 6D, F). Using RT-qPCR, the upregulation of these genes by MDA231-HEXOs were confirmed significant except for three genes in the EMT gene set (COL4A1, ITGAV, TPM1) and seven genes in the OP gene set (SUCLA1, ETFA, COX6A1, TIMM8B, UQCRQ, ATP5PF, NDUFA4) (Fig. 6D, F). Interestingly, almost all of these mRNAs were carried by the MDA231-HEXOs except for two collagen genes (COL5A2, COL4A1) and the SUCLA1 mRNA which were also absent in HUVECs (Fig. 6D, F). Therefore, in addition to the hypoxia-targeted mRNAs, the MDA231-HEXOs also carried the metabolism-related and motility-related mRNAs, which at least partially contributed to the upregulation of these genes in HUVECs.

## Discussion

Exosome cargoes have been extensively investigated to discover biomarkers for cancer diagnosis. Among the exosomal molecules, proteins and miRNAs are currently the two major types that have been widely investigated for cancer diagnosis. Several exosome databases have been established to collect extensive exosomal proteins and miRNAs^40-44^. However, the feasibility of exosomal mRNAs as biomarker is still not well studied. Here, we quantified 61 hypoxia-targeted mRNAs in exosomes and found that 30 of these mRNAs were present in cancer exosomes and absent in normal exosomes. Therefore, we proposed that the unappreciated exosomal mRNA is a promising source of cancer biomarker. Recently in liquid biopsy for cancer diagnosis, cell-free DNA (cfDNA) is becoming a leading type of biomarker alongside the technological advances in nucleic acid quantification technologies, such as PCR and next generation sequencing (NGS)^45^. Nevertheless, using cfDNAs as biomarkers faces challenges. For example, the molecular origin of cfDNAs remains controversial, and the most likely sources are dead cells which passively release cfDNAs. Furthermore, the cfDNAs are exposed to the DNase in body fluids and thus prone to degradation. As a result, the level of cfDNAs may randomly fluctuate, rendering such information less reproducible. In contrast, exosomes are actively released through endosomal pathway, with the cargo well-protected by the enclosed membrane^15^. Notably, we found the exosomal mRNAs levels tend to be homeostatic since the coefficient of variance of independent replicates were all small (below 0.3). Other works also suggest that the levels of exosomal mRNAs may reflect the physiological status of original cells^17,46^. As both exosomal mRNAs and cfDNAs can be quantified using the advanced nucleic acid quantification technologies with high sensitivity, we envisaged that exosomal mRNA is a competitive alternative to cfDNA, mainly because the actively secreted and membrane-enclosed exosomal mRNAs are thought to be more informative, stable and abundant.

Among the identified mRNAs that were present in cancer-derived exosomes and absent in MCF10A-derived exosomes, several mRNAs including SNAIL1/2, TAZ, AXL, RHOA and ROCK1 are well known to promote cell motility. SNAIL1/2 and TAZ are transcriptional factors or co-activators that induce EMT by suppressing epithelial markers like E-cadherin and upregulating mesenchymal markers including vimentin, fibronectin as well as DOCK4/5/9^47-50^. AXL receptor tyrosine kinase demonstrates an exacerbating feedback with SNAIL1/2 that the overexpression of AXL elevates the SNAIL1/2 expression, and vice versa^51,52^. RHOA/ROCK signaling axis can mediate the formation of actin stress fibers and focal adhesions, which can enhance the lamellipodium-based cell migration^53^. Taken together, these previously unappreciated pro-migratory mRNAs may synergize with the exosomal proteins like MET^54^, AREG^55^, TGFβ1^56^, Wnt11^57^ and HIF1α^20^ as well as exosomal miRNAs like miR-21^27^ to educate the recipient cell towards a migratory phenotype. Specifically for HUVECs, we observed an upregulation of EMT gene set by the treatment of MDA-MB-231 derived exosomes. The endothelial-mesenchymal transition of HUVECs can lead to the formation of cancer-associated fibroblast^58^, the increase of angiogenic sprouting^59^, and the elevation of blood vessel permeability^60^, all of which contribute to the tumor progression. The integrity of the exosomal mRNAs has been addressed by several works but it remains controversial. RNA size profiling by Agilent bioanalyzer shows short length RNAs are enriched in exosomes^3^. Also, microarray analysis indicates that 68.5% exosomal mRNAs are likely to be in fragments^61^. These results suggest full-length mRNA may be far less abundant than short length RNAs in exosomes. Nevertheless, the fragmentation of exosomal mRNAs does not necessarily prevent them from being biomarkers. A previous study demonstrated that detecting the short length segment (∼200bp) of EGFR mutant in exosomal nucleic acids exhibits higher sensitivity (76.5%) than cfDNAs (64.7%) in diagnosing NSCLC patient^62^. Meanwhile, evidence suggest at least some full-length translatable mRNAs are present in exosomes. For example, mice cell MC/9 derived exosomal RNAs are translated into proteins by *in vitro* rabbit lysate translation kit, and the seven translated proteins are confirmed to be of mouse origin^3^. In addition, the exosomes secreted by glioblastoma cell transduced with Gluc transfer Gluc mRNA to HBMVEC cells and induce continuous increase of Gluc activity over 24 h^15^. We also observed that MDA231 cells transfected with GFP-tagged VEGFA derived exosomes that carried GFP mRNA. The GFP-tagged VEGFA were transferred to HUVECs and translated to fluorescent GFP protein in 24 hours. Taken together, the translatable mRNAs are present in exosomes and may produce noticeable levels of proteins in recipient cells regardless of their relatively low abundance in exosomes.

The comparative characterization of exosomes from MCF10A, MCF7 and MDA231 cell revealed that cancer cells transferred prominently more molecules via exosomes than normal cells. With the quantitative characterization of the exosomes, we noticed the rate of exosome secretion by exocytosis positively correlated with cell motility among MCF10A, MCF7 and MDA231 cell lines. This observation is supportive by a cell migration theory - the membrane flow model - whereby the exocytosis together with endocytosis and vesicle trafficking constitute a cycle of rearward membrane flow which contributed to cell migration^63-66^. The speed of the membrane flow, which represents the frequency of exocytosis, is found higher in cells migrating with faster velocity^65^. Therefore, the cellular activities of exosome generation, exocytosis, membrane flow and cell migration are likely to be associated. The increased frequency of exosome secretion by migratory cancer cells may serve to elevate the level of circulating exosomes in cancer patient, supplementing the previous understanding that the increased secretion of exosomes is induced by the stresses in tumor microenvironment like hypoxia^67-69^. Furthermore, we observed that MDA231 cells not only secrete more exosomes, but also produced larger exosomes than MCF10A and MCF7 cells. These results indicate that besides exosome cargoes, the biophysical features of exosomes may also be informative for cancer diagnosis. Additionally, we found remarkably more types of mRNAs were present in cancer exosomes than normal exosomes. Therefore, cancer cells loaded more types of molecules into the exosomes and secreted the exosomes more frequently than normal cells.

Hypoxia as a typical stress in the TME induces the formation of new blood vessels, i.e. angiogenesis, which in turn boosts the supply of oxygen to mitigate the hypoxic stress. This feedback response is achieved through the key mediator, VEGFA. First, hypoxia promotes the cytoplasmic level of HIF-1 which binds to a 47-bp hypoxia response element in VEGFA transcription initiation site to upregulate the VEGFA expression^70^. Subsequently, the elevated VEGFA initiates angiogenesis by binding to the VEGF receptors present on the surface of endothelial cells, which activates downstream signaling pathways such as MAPK/ERK pathway^71^. In this study, we found that high level of VEGFA mRNA was carried by exosomes derived from multiple types of cancer, and the level was modulated by the hypoxic stimulus. Furthermore, treating HUVECs with the VEGFA mRNA^+^ exosomes promoted angiogenesis via VEGFR-dependent pathway. These results indicate that cancer cells may favor angiogenesis not only through secretory VEGFA protein, but also by transferring VEGFA mRNAs via exosomes.

Study of hypoxia-induced cancer exosomes is still at its stage of infancy. Evidences suggest that hypoxia-induced cancer exosomes contribute towards transforming the neighboring cells and resulting in angiogenesis, invasion, metastasis and immune evasion. These effects have been mainly attributed to specific proteins and miRNAs derived from the exosomes. For example, exosomal proteins (TF^22^, Wnt4^72^) and miRNAs (miR-135b^28^, miR-23a^21,26^, miR-210^73^, miR-494^74^) are upregulated by hypoxia and thus advance angiogenesis. Similarly, hypoxia increases the exosomal miR-21^27^ and lncRNA-UCA1^75^ which promotes EMT. In line with these results, we observed that hypoxia-induced MDA231-HEXOs promoted the VEGFR-dependent angiogenesis and elevated the expression of EMT-related genes in HUVECs. Rather than proteins and non-coding RNAs, the identification of angiogenesis-related mRNAs and EMT-related mRNAs in the exosomes indicates that the mRNAs are at least partially implicated in reprograming the recipient cells. Interestingly, we found that the exosomal mRNAs derived from the two cancer cells MCF7 and MDA231 responded to hypoxia in an opposite manner. With hypoxic stress, most of the hypoxia-targeted mRNAs were downregulated in MCF7-EXOs but upregulated in MDA231-EXOs. This contradicting response to hypoxia could not be adequately explained with the HIFs-dependent transactivation of genes which always upregulates the target genes. Future investigation of the machinery that packs mRNAs into exosomes could extend our understanding of how the hypoxia regulates the exosomal mRNAs, thus opening up potentially new revenues for diagnosis and treatment of cancer.

## Methods

### Cell culture

MCF10A, MCF7, MDA-MB-231, Caco2, SW480, SW620, MIAPaCa-2, HepG2 were purchased from American Type Culture Collection (ATCC, Manassas, VA, USA). H1650 and H1792 were obtained from National Cancer Center Singapore. GC38 was a gift from Dr. Shing-Leng Chan from Cancer Science Institute of Singapore. As for the culture medium, the cells MCF7, MDA-MB-231, SW480, SW620, MIAPaCa-2 and GC38 were cultured in DMEM (Thermo Fisher Scientific, Cat No. 11965092) supplemented with 10% Fetal Bovine Serum (FBS; Thermo Fisher Scientific, Cat No. 26140079). Caco2 was cultured in DMEM supplemented with 20% FBS. HepG2 was cultured in EMEM (Thermo Fisher Scientific, Cat No. 11095080) supplemented with 10% FBS. H1650 and H1792 were cultured in RPMI-1640 (Thermo Fisher Scientific, Cat No. 11875093) supplemented with 10% FBS. These cells were cultured under 20% O2 for normoxia and 1% O2 for hypoxia.

### Exosome incubation and isolation

Confluent cells in T75 cell culture flask were starved in serum-free medium for one day in either normoxia or hypoxia incubator. Then the medium was replaced with fresh serum-free medium, and the cells were incubated for another 24 hours to produce exosomes. The exosomes in culture medium were enriched and purified by differential centrifugation following the established protocol^29^. Briefly, cell debris was removed by 10 min and 2,000 x g centrifugation. Larger vesicles including apoptotic bodies and microvesicles were excluded by 30 min and 10,000 x g centrifugation. Finally, exosomes were pelleted by 2 h and 100,000 x g centrifugation and washed by PBS. The enriched exosomes were kept in 4 °C and processed in less than 3 days or stored in negative 80 °C for long-time storage.

### Quantitative analysis of exosomes by Nanoparticle Tracking analysis

A number of MCF10A, MCF7 and MDA-MB-231 cells were cultured in T25 flasks. Cells were first starved in serum-free medium for 3-6 hours, followed by PBS wash for three times and the refresh of the serum-free medium. After 3 h incubation, the medium was collected, and the larger vesicles in culture medium were removed by 30 min 10,000 x g centrifugation. The remaining medium containing exosomes was diluted properly and analyzed by the NanoSight LM10 (Malvern Panalytical, Malvern, UK). Meanwhile, the cells in the T25 flask were harvested and counted by the Multisizer 4e coulter counter (Beckman Coulter, Brea, CA, USA).

The accuracy of the quantitative analysis were optimized with the following measures: first, the exosome incubation is constrained to a short 3-hour period that is much shorter than the cell doubling time (more than 16 h) so that the total cell number is minimally affected by cell proliferation; second, the samples with discrepant concentrations were diluted accordingly to achieve similar final concentration, thus the comparison of the concentrations between samples became reliable independent of the technical linear range of the NTA; third, the data of each sample was acquired by averaging the measurements of 10 random regions, reducing the effects of the heterogeneity among regions.

### PKH26 staining of cells and exosomes

Confluent cells were trypsinized and stained by PKH26 Red Fluorescent Cell Linker Kit (Sigma-Aldrich, Cat No. PKH26GL) following the vendor’s instruction. Briefly, less than 1×10^7^ cells were pelleted and resuspended in 1 mL Diluent C, and then mixed with 1 mL 4×10^−6^ M PKH26 for 5 min. Next, the staining was stopped by incubating with 2 mL fetal bovine serum for 1 min. Then the stained cells were wash with complement culture medium for three times to remove excessive dye. Last, the cells were seeded in flasks to produce exosomes, or in petri dishes for fluorescent imaging.

To stain the exosomes, we refrained from directly mixing the PKH26 dye with exosomes because the excessive dye co-isolated with exosomes and mimicked exosomes even in control PBS sample (Fig. S1A). Such issue of contamination by excessive PKH dye was also reported in other study^76,77^. Instead, the PKH26 dye was used to only stain the cells, followed by triple wash with complete culture medium to thoroughly remove the excessive dye. Taking the advantages that the PKH26 staining is efficient and the fluorescence last for weeks, the exosomes generated by the PKH26^+^ cells were also positive with PKH26 (Fig. S1B).

### Western blot

The proteins of cells and exosomes were extracted with the RIPA as described in the previous study^78^. The total protein levels of the lysates were measured by BCA protein assay kit (iNtRON Biotechnology, Cat No. 21071). The following primary antibodies were used at 1:1000 dilution: Alix (Cell Signaling Technology, Cat No. 2171), CD63 (Abcam, Cat No. ab59479), CD9 (Abcam, Cat No. ab92726), CD81 (Abcam, Cat No. ab79559), VEGFA (Abcam, Cat No. ab214424), VEGFR1 (Abcam, Cat No. ab32152), VEGFR2 (Abcam, Cat No. ab134191), Erk1/2 (Cell Signaling Technology, Cat No. 4695), β-actin (Thermo Fisher Scientific, Cat No. MA5-15739-HRP). The primary antibody phospho-Erk1/2 (Sigma-Aldrich, Cat No. M9692) was used at 1:3000 dilution. The secondary antibodies were used at 1:3000 dilution.

### RT-qPCR and DNA gel electrophoresis

Total cell RNA was extracted by RNeasy Kit (Qiagen, Cat No. 74106). Total exosome RNA was isolated by Total Exosome RNA & Protein Isolation Kit (Thermo Fisher Scientific, Cat No. 4478545). RNA was revere transcribed to cDNA by SuperScript III Reverse Transcriptase (Thermo Fisher Scientific, Cat No. 18080044), followed by RT-qPCR with the FastStart Universal SYBR Green Master (Sigma-Aldrich, Cat No. 4913850001). Primers are listed in Supplementary Table 2. Two percent agarose gel was prepared by dissolving 2 g Agarose (Bio-rad, Cat No. 1613102) and 2 µL SYBR Safe DNA Gel Stain (Invitrogen, Cat No. S33102) in 100 mL Tris-acetate-EDTA (TAE) buffer. Amplified DNA of target gene was mixed with 6x DNA gel loading dye (Thermo Fisher Scientific, Cat No. R0611) and subjected to gel electrophoresis at 105 V for 30 min. GeneRuler 100 bp DNA Ladder (Thermo Fisher Scientific, Cat No. SM0241) was used as the reference of DNA size. Imaging of the gel was done by iBright FL1000 (Invitrogen).

### Confocal imaging of cells and exosomes

Cells were fixed on glass by 4% paraformaldehyde solution in PBS. Hoechst 33342 (Thermo Fisher Scientific, Cat No. 62249) was used at 1:1000 dilution to stain cell nucleus. The cell actin was stained by Phalloidin-iFluor 647 (Abcam, Cat No. ab176759) at 1:2000 dilution for 30 min. After the staining, the fixed sample were imaged by Nikon A1R confocal microscopy.

Exosomes were sealed on glass slides for imaging. Briefly, 10 µL exosomes were dripped on the glass slide. Coverslip was placed on top of the drop to spread the liquid. Next, nail polish was applied to immobilize and seal the coverslip. The slide with the exosomes was imaged by Nikon A1R confocal microscopy with the coverslip side facing the lens.

### Imaging of exosome transfer

CD63-GFP was a gift from Paul Luzio (Addgene plasmid # 62964). MDA-MB-231 cells were transfected with the CD63-GFP by Lipofectamine 3000 (Thermo Fisher Scientific, Cat No. L3000015). The CD63-GFP^+^ MDA-MB-231 cell were selected by gentamicin and enriched by flow cytometry. One day before co-culture, 1×10^4^ HUVECs were seeded in glass-bottom 8 well µ-slide and incubated overnight. After the HUVECs adhered, 1×10^4^ GFP^+^ MDA-MB-231 were harvested, stained with PKH26 and added to the HUVECs. After 24-hour co-culture in 1:1 mixed EGM-2 and FBS-supplemented DMEM, the cells were fixed and imaged at 100x magnification by the confocal Nikon A1R.

### Live cell imaging of RNA transfer

About 1×10^5^ MDA-MB-231 cells were stained by PKH26 and seeded in 35mm glass bottom petri dish. After the MDA-MB-231 cells adhered, the RNA of MDA-MB-231 was stained by SYTO RNASelect green fluorescent cell stain (Thermo Fisher Scientific, Cat No. S32703) at 1:10,000 dilution. Excessive dye was removed by three times PBS wash. Next, 1×10^5^ HUVECs were harvested and added to the PKH26^+^ and SYTO RNASelect^+^ MDA-MB-231. The petri dish was put in live cell imaging chamber immediately and imaged at 60x magnification continuously with 1 h interval.

### Exosomal transfer of VEGF-GFP mRNA

The VEGFA-GFP plasmid was constructed by the protein cloning Core of Mechanobiology Institute in Singapore. The GFP were fused with VEGF165 at the c-terminus of VEGF165. MDA-MB-231 cells were first stained with PKH26, followed by transient transfection of the VEGFA-GFP plasmid with Lipofectamine 3000 (Thermo Fisher Scientific, Cat No. L3000015). The next day, the transfected cells were incubated in serum-free DMEM for exosome generation. After 24 h, the culture medium was collected, and the exosomes in the culture medium were enriched and purified. These exosomes were then used to treat the HUVECs for 24 h. Last, the HUVECs were fixed and imaged with 100x magnification.

### Tube formation assay

The 24 well plate was coated with 80 µL growth factor reduced Matrigel (Bio Laboratories, Cat No. 354263). After gel formation, 1×10^5^ HUVECs were seeded in each well. The cells were incubated in EBM-2 basal medium for control, or treated with 50 µg/mL MDA-MB-231-HEXOs, or treated with both 50 µg/mL MDA-MB-231-HEXOs and 0.1 µM Axitinib (Selleck Chemicals, Cat No. S1005). After 24 h, the cells were stained with Calcein AM (Thermo Fisher Scientific, Cat No. L3224), following by imaging of multi positions under 10x magnification. Tube length in each image was analyzed by the Angiogenesis Analyzer plugin for ImageJ^79^.

### EdU Proliferation assay

The Click-iT™ EdU Proliferation Assay kit (Thermo Fisher Scientific, Cat No. C10499) was used for the proliferation assay of HUVECs following the manufacturer’s instruction. Briefly, HUVECs were seeded in 24 well plated and incubated for overnight. The adhered HUVECs were starved in EBM-2 for 3 hours, followed by incubation with 10 µM EdU in either basal EBM-2 or 50 µg/mL MDA-MB-231-HEXOs. After 24 hours, the HUVECs were fixed by Click-iT EdU fixative. Then the cell nucleus was stained by Hoechst 33342. By fluorescent imaging, the total cells were counted with nucleus fluorescence and the proliferative cells were counted with EdU fluorescence.

### RNA-seq and data analysis

HUVECs were incubated in basal EBM-2 for control or treated with 50 µg/mL MDA-MB-231-HEXOs for 24 h. The total cell RNA of two control samples and two treated samples were extracted with RNeasy Kit (Qiagen, Cat No. 74106), and then processed and sequenced by BGI Genomics (BGI Genomics, Shenzhen, China). With the RNA-seq data, the sequencing quality were verified and the correlation between samples were analyzed by BGI Genomics. The filtered clean reads were mapped to the human genome reference GRCh38.p13 Ensemble release 97 by two parallel pipelines, HTSeq^80^ and Salmon^81^. Based on the gene counts acquired by the two algorithms, differentially expressed gene (DEGs) were analyzed by the two competing approaches DESeq2^82^ and edgeR^83^ with the thresholds of 0.05 adjusted p-value or 0.1 False discovery rate, respectively. As a consequence, four lists of DEGs were acquired and a final DEG list was obtained by intersecting the four lists. The gene ontology enrichment analysis of the DEGs was completed on The Gene Ontology Resource (http://geneontology.org/). The gene set enrichment analysis (GSEA) was finished with the GSEA software^84^.

## Data availability

The data that support the findings of this study are available from the corresponding author upon reasonable request.

## Acknowledgements

This work was carried out at the MechanoBioEngineering (MBE) laboratory at the Department of Biomedical Engineering, National University of Singapore (NUS) and the Mechanobiology Institute (MBI), NUS. We thank the members of the MBE lab for their support. We thank Paramasivam Kathirvel from MBI for constructing the VEGFA-GFP plasmid. P. Z. gratefully acknowledges scholarship and support from NUS Graduate School for Integrative Sciences and Engineering (NGS), NUS.

## Author contributions

P.Z., B.C.L. and C.T.L. planned the study. P.Z. and S. B. L. performed mRNA quantification. P. Z. and K. J. performed cell imaging. P. Z. and T. W. C performed western blotting. P. Z., B. C. L. and C. T. L. analyzed and interpreted the data. P. Z., S. B. L., K. J., T. W. C., B. C. L. and C. T. L. wrote and reviewed the manuscript.

## Figures

**Fig. S1:**
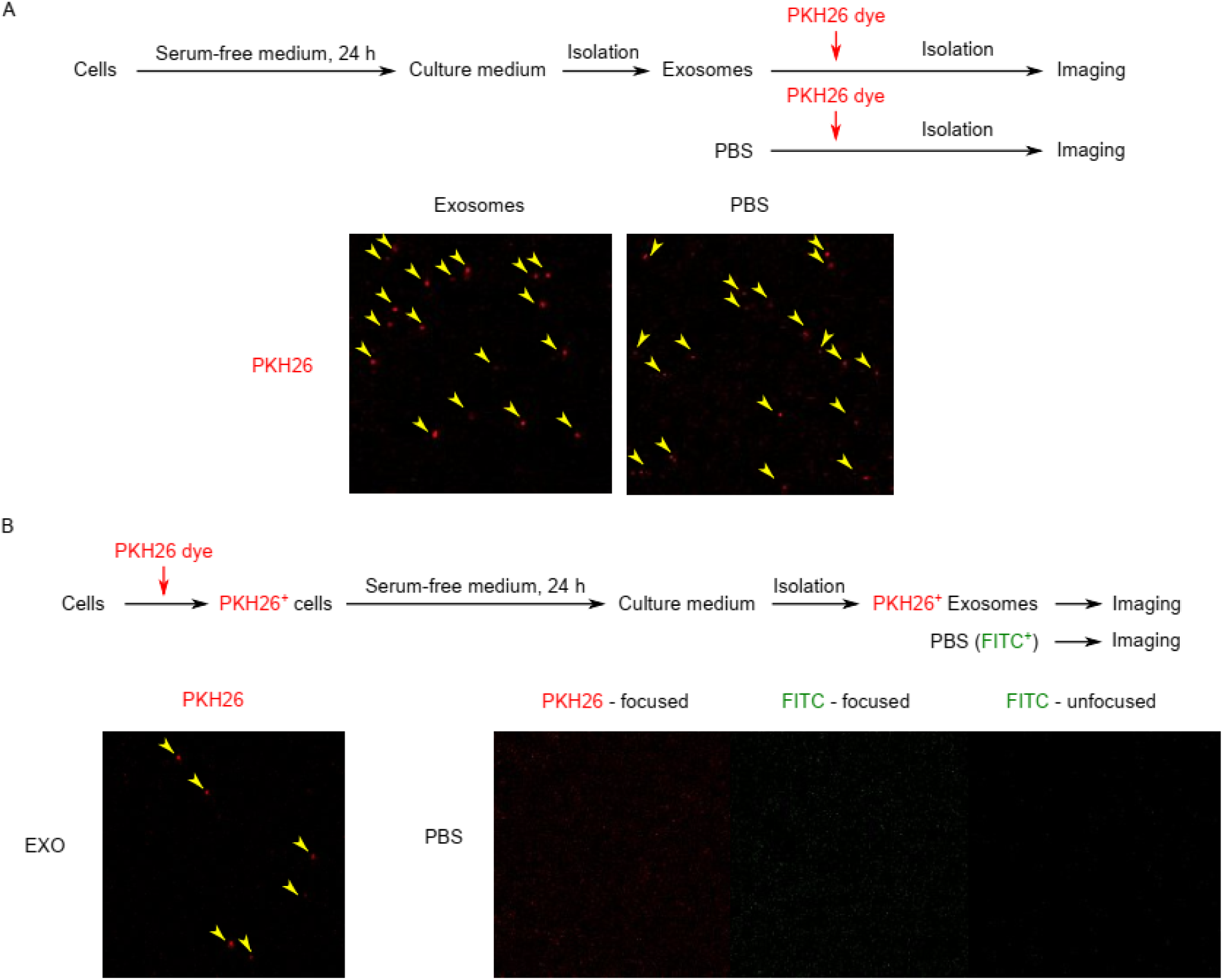
Comparison of two methods of exosomes staining with PKH26. (A) Staining of exosomes with direct mixing of exosomes and PKH26 dye introduced excessive dye that co-isolated with exosomes and mimicked exosomes in fluorescent imaging since abundant exosome-like spots were identified in PBS control sample. (B) Cells that were stained with PKH26 dye generated PKH26^+^ exosomes. Excessive dye was thoroughly wash off after cell staining. PBS control showed no fluorescence.

**Fig. S2:**
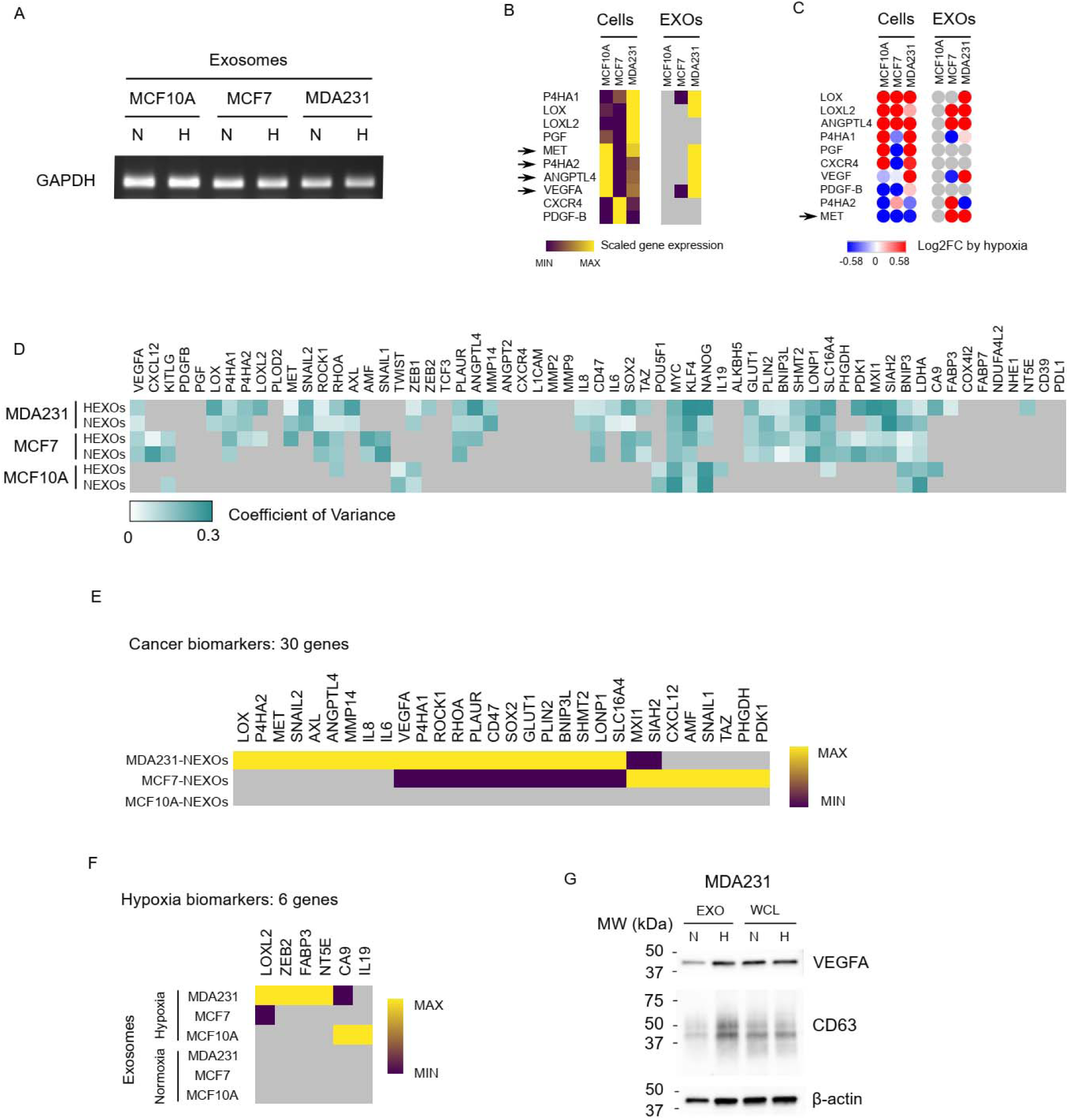
Quantification of hypoxia-targeted mRNAs in exosomes. (A) GAPDH mRNA was robustly present in all the quantified exosomes. (B) Comparison of gene expressions in cells and exosomes. The expression levels were normalized to GAPDH. (C) Comparison of hypoxia induced alterations of mRNA level in cells and exosomes. (D) The coefficient of variances of all gene expressions were less than 0.3, demonstrating that the exosomal mRNA levels were homeostatic, and the quantification of exosomal mRNA by RT-qPCR was reproducible. (E)(F) Heatmap shows the exosomal mRNA level of the 30 genes recognized as cancer biomarkers and the 6 genes regarded as hypoxia biomarkers. (G) Western blot shows that the VEGFA protein level in cells (normalized to β-actin) and in exosomes (normalized to CD63) were not responsive to the hypoxic stimulus.

**Fig. S3:**
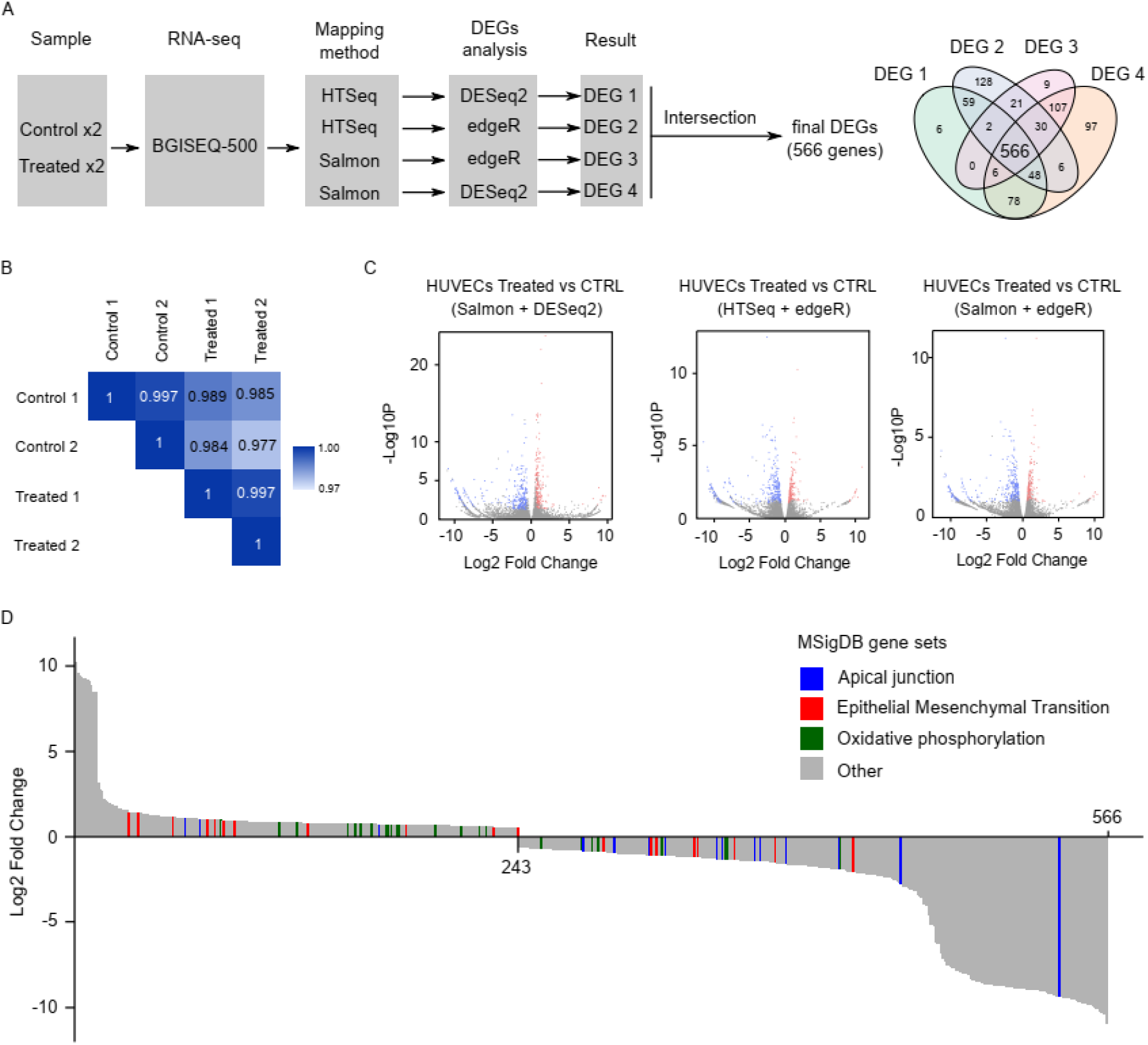
RNA-seq of HUVECs treated with cancer exosomes. (A) Workflow of the RNA-seq data analysis. The final list of DEGs comprising 566 genes were acquired from intersection of four list of DEGs. (B) Correlation of four samples based on the expression profile. (C) Volcano plot shows RNA-seq data of HUVECs with or without the treatment of MDA231-HEXOs that were analyzed with Salmon + DESeq2, HTSeq + edgeR, or Salmon + edgeR packages. (D) The DEGs ranked with the fold change. Genes that belongs to Apical junction gene set, EMT gene set, or OP gene set were highlighted in blue, red, or green color, respectively.

